# Impacts of heminode disruption on auditory processing of noisy sound stimuli

**DOI:** 10.64898/2026.02.02.703242

**Authors:** Siddhant Tripathy, Maral Budak, Ross Maddox, Anahita H. Mehta, Michael T. Roberts, Gabriel Corfas, Victoria Booth, Michal Zochowski

**Author notes:** (MZ); (VB).

## Abstract

Hidden hearing loss (HHL) is an auditory neuropathy characterized by altered auditory nerve responses despite normal hearing thresholds. Recent experimental and computational studies suggest that permanent disruptions to heminode positions in spiral ganglion neuron (SGN) fibers can contribute to these deficits. However, the interaction between heminode disruption and noisy backgrounds ubiquitous in daily listening remains unexplored. This study investigates how background noise affects auditory processing with these peripheral disorders and how deficits propagate to downstream sound localization circuits in the superior olivary complex. We developed computational models of SGN fibers with mild and severe degrees of heminode disruption, subjected to sinusoidal tone stimuli in the presence of background noise with varying spectral characteristics. We analyzed the phase-locking of SGN fiber responses to the stimulus tone and modeled the subsequent effects on interaural time difference (ITD) sensitivity in the medial superior olive (MSO) using a binaural localization network. We found that near-tone-frequency noise disrupted SGN phase locking through cycle-to-cycle variability in spike phases, with effects consistent across tone frequencies. Mild heminode disruption produced frequency-dependent degradation in SGN phase locking, with effects observed only at higher frequencies tested (600–1000 Hz), without reducing overall firing rates. Critically, the effects of noise and heminode disruption were additive, with combined exposure leading to reduced ITD sensitivity and large temporal fluctuations in MSO responses. Severe heminode disruption, which additionally reduced firing rates at the SGN fibers and subsequent stages, produced profound localization deficits across all frequencies tested. Thus, our model results suggest that noisy environments exacerbate auditory deficits from peripheral disorders implicated in HHL and could potentially impair speech intelligibility through degradation in localization ability. This model may be useful for understanding the downstream impacts of SGN neuropathies.

## Introduction

Hidden hearing loss (HHL) is an auditory neuropathy characterized by changes in sound-evoked neural output of the auditory nerve (AN) without hearing threshold elevation [1]. In animal studies, signatures of HHL can be detected directly as an attenuation of the auditory brainstem response (ABR), a far-field response measured by head-mounted electrodes, whose first peak (ABR peak 1) reflects the cumulative activity of auditory nerve fibers [1], specifically type 1 spiral ganglion neurons (SGNs, Fig 1A).

**Fig 1.**
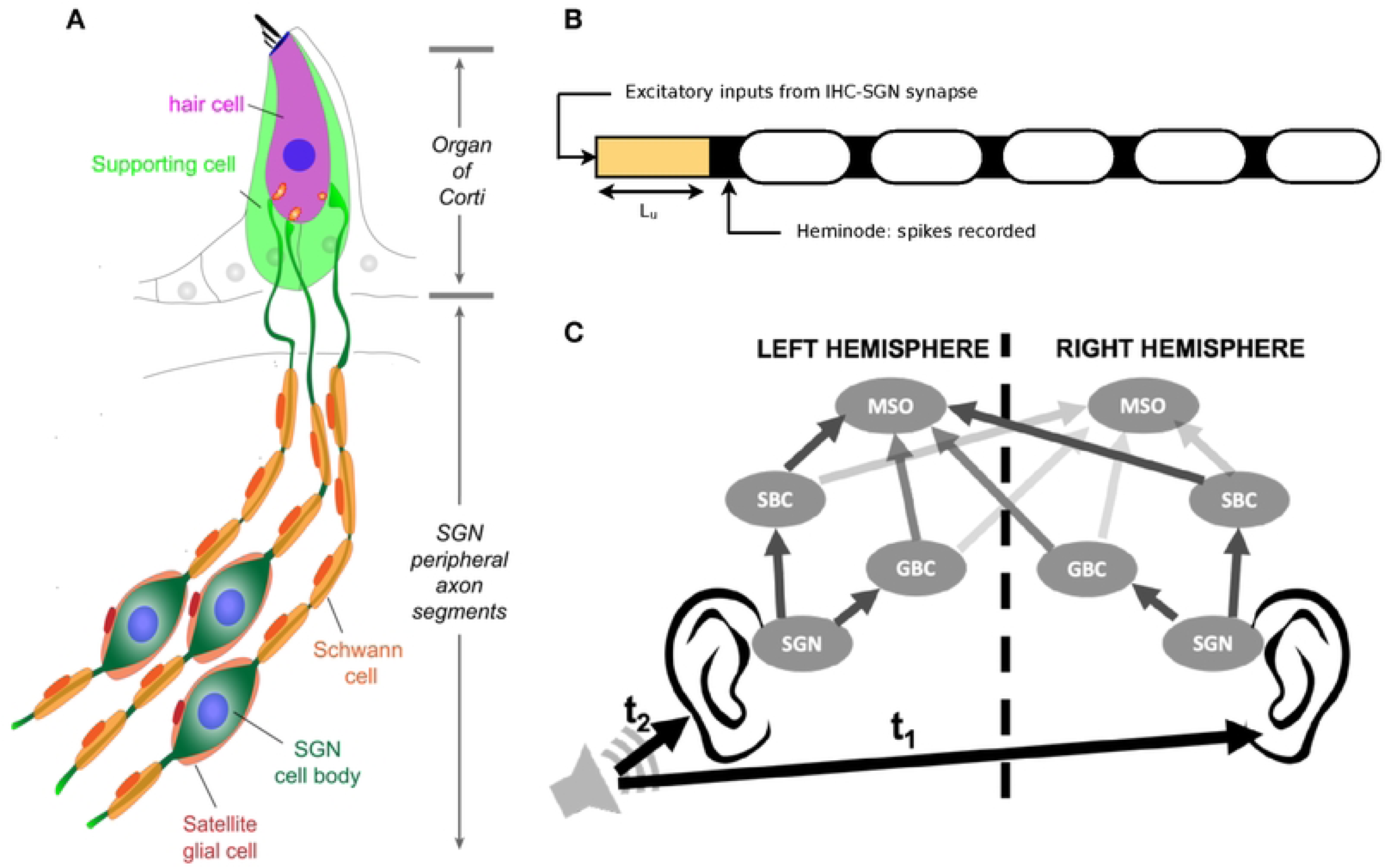
The auditory processing model. A: Schematic illustration of type I Spiral Ganglion Neurons (SGNs) innervating Inner Hair Cells (IHCs) via myelinated peripheral projections (adapted from Figure 1 in [12, 13]). B: Schematic of our axon cable model implemented with python package NEURON, consisting of an initial unmyelinated segment of length L_u_ adjacent to the site of IHC synapses, followed by a heminode and 5 myelinated segments with 4 interconnecting nodes. Neurotransmitter release at IHC synapses was modeled as excitatory inputs to the SGN fiber at the beginning of the unmyelinated segment, and the action potentials (APs) at the heminode were recorded for all fibers simulated. C: Cochlear nucleus circuit model for studying localization of sound sources in the azimuthal plane, consisting of SGNs, Spherical Bushy Cells (SBCs), Globular Bushy Cells (GBCs), and Medial Superior Olives (MSOs) on both sides (adapted from Figure 1 in [14]). SGN spikes are relayed to SBCs and GBCs on the ipsilateral side only, which in turn relay their inputs to MSOs on each side, both ipsilaterally and contralaterally. SBC inputs to the MSO are excitatory, while GBC inputs are inhibitory. The azimuthal location of the sound source is encoded in the Interaural Time Difference (ITD), the difference in the arrival time of the sound to both ears (t_1_-t_2_). The difference in the spike rate between the left and right MSOs is sensitive to the ITD and was studied as a measure of the localization performance of the circuit.

In humans, HHL has not been formally shown, with mixed reports of presumptive auditory nerve damage resulting in perceptual deficits without hearing threshold changes [2–5]. It has been hypothesized that a reduction in the ability to localize sound sources in the horizontal plane could contribute to such perceptual deficits. This localization takes place in the superior olivary complex (SOC), specifically in the medial superior olive (MSO), and involves the precise timing of auditory signals coming from both ears in response to a sound stimulus. In particular, the MSO integrates binaural inputs from ipsilateral and contralateral populations of bushy cells in the cochlear nucleus [6]. Each bushy cell in turn receives input from multiple SGNs and is tuned to integrate these signals into precise phase information that is transmitted to the MSO. Thus, pathologies that disrupt the signaling along peripheral auditory circuits could impair downstream information processing by disrupting the timing of inputs arriving at each stage of the auditory pathway, resulting in degraded sound localization ability, potentially affecting speech intelligibility [7].

These pathologies can be associated with the loss of synapses between inner hair cells (IHCs) and SGN fibers (Fig 1A), even if the complement of hair cells and SGNs remains unchanged [8]. This type of pathology, termed synaptopathy, has been implicated in HHL and can occur in response to prolonged exposure to loud noise but also in healthy aging [9, 10].

Since proper myelination of auditory nerves is essential for auditory processing [11], myelin disorders could also cause peripheral neuropathies that lead to HHL. Recent studies in mice showed that acute demyelination of SGN fibers resulted in decreased ABR peak 1 amplitudes, and additionally, increased ABR peak 1 latencies, without auditory threshold elevation or loss of IHC-SGN synapses [12]. Remarkably, these changes persisted even after complete remyelination of SGN fibers. Further immunostaining experiments showed disruptions in the positions of heminodes, the first nodes on SGN fiber axons, closest to the IHCs, where action potentials are generated in response to incoming sound stimuli.

By incorporating variability in heminode locations in SGN fibers, our previous computational modeling efforts successfully replicated the ABR peak changes observed experimentally [13]. Modeling results suggested a two-fold effect of heminode disorganization in SGN fibers on the ABR: lower SGN activity levels and asynchronous activity across SGN fibers. As a result, small amounts of jitter in SGN activity in response to sound input were greatly amplified by this disorder, thereby impairing the convergence of inputs for subsequent processing. We note here that, while we have referred to such heminode disruption as myelinopathy in our previous work, the term can generally refer to a broader class of myelin disorders. In this study as well, we continue to use the term myelinopathy to refer to our modeling of heminode disruption specifically. Our previous efforts also compared the effects of myelinopathy (specifically heminode disorganization) and synaptopathy [13], and showed that synaptopathy reduced SGN activity levels without affecting response timing. In an extended network model of the peripheral system and cochlear nuclei and MSO populations, we found that the effects of both pathologies at the peripheral level significantly reduced the sensitivity of MSO responses to the horizontal location of brief sound inputs, which could potentially explain some perceptual deficits in humans [7, 14].

In realistic scenarios, a sound source is accompanied by background noise of varying spectral content, which has been shown to introduce perceptual difficulties even in healthy hearing conditions [15]. With background noise present, the sound waveform additionally contains several frequency components at random phases, which may affect the phase locking of SGN fibers to the stimulus signal. How such noise-induced jitter in SGN firing activity interacts with myelinopathy deficits to affect localization processing in the MSO has not been determined.

In this study, we used computational models for the peripheral auditory system, cochlear nuclei, and the MSO to investigate the impacts of myelinopathy on auditory processing and sound localization in the presence of background noise. We focused on sound frequencies between 200 Hz and 1000 Hz, a frequency regime in which SGN fibers show robust phase locking with sound waveforms [16]. With a spike-time-based model, we analyzed perturbations to the fine temporal structure of individual SGN fiber spiking responses to sound stimuli, instead of considering ABRs that capture population-level SGN activity for a short stimulus window [16–18]. Thus, we identified how noise and myelinopathy together affect SGN spiking patterns in distinct ways that lead to auditory processing deficits. In contrast to our previous work, here we considered two regimes for myelinopathy: a mild regime where there is no decrease in overall SGN firing activity, allowing us to isolate only the effects of temporal disruptions, and a severe regime that additionally accounts for reduced SGN activity. We then analyzed how changes at the SGN level impact downstream localization processing in the MSO in noisy environments. Model results showed that noise frequencies close to the frequency content of the sound stimulus degraded the phase locking of SGN fibers tuned to those frequencies, and this degradation was more severe with even mild myelinopathy. Consequently, mild myelinopathy led to a reduction in localization ability despite healthy levels of activation of SGN fibers. Additionally, in the MSO, background noise caused random fluctuations in the sensitivity to stimulus location throughout the duration of a sound stimulus. In the severe myelinopathy regime, localization deficits were severely exacerbated due to reduced levels of activation of SGN fibers, and consequently of cochlear nuclei and MSO cells. Taken together, our model results suggest that peripheral deficits caused by myelinopathy in noisy conditions lead to a degradation in localization ability that could potentially explain some of the speech intelligibility deficits reported in conditions like the ‘cocktail party problem’ in humans [7, 19].

## Materials and methods

To analyze the impacts of peripheral deficits implicated in HHL on auditory processing, we used our previously developed physiology-based computational model that simulates neural activity along the auditory pathway in response to a given input signal waveform [13, 14], with some modifications where needed. The model consists of a peripheral auditory processing model to generate synaptic inputs to auditory nerve fiber models in response to sound with noisy background (Fig 1B). Auditory nerve fiber activity is input to Superior Olivary Complex (SOC) models that include neural populations up to the Medial Superior Olive (MSO), where the horizontal localization of sound is processed (Fig 1C). While the details of the equations and all parameters will not be repeated here (see [13, 14]), we will briefly describe the model and emphasize any changes we have made.

### Sound inputs

We constructed input sound waveforms consisting of the linear superposition of a pure tone stimulus and background noise. The tone was chosen to be a pure sinusoidal tone of frequency 200 Hz, 600 Hz, or 1000 Hz, covering a range of frequencies where the auditory system shows robust phase locking with the fine temporal structure of sound stimuli [16]. Each sound waveform was 100 ms long, comprising 40 ms of background noise only, followed by 60 ms of the tone superimposed on the background noise. For the background, we considered 5 cases: a case with no noise which serves as the control (Fig 2A shows an example for a 600 Hz tone), a white noise case, and 3 narrowband noise conditions: one with the center frequency of the noise spectrum at the tone frequency (called centered noise, Fig 2B), one with center below and one with center above (Fig 2C) the tone frequency. The power spectra for the sound stimuli comprising the pure tone superposed with each of these 5 background cases are shown in Fig 2G-I for all 3 tone frequencies. The narrowband noise waveforms were created by applying a second-order Butterworth bandpass filter to additive white Gaussian noise (AWGN), with a width of 80 Hz for the tone frequency at 200 Hz, and a width of 150 Hz for 600 Hz and 1000 Hz tones. For 600 Hz and 1000 Hz tones, the sound pressure level (SPL) of the tone signal was chosen to be 66 dB, a reasonable estimate for SPL levels of human conversation and animal sounds [20–22]. For the 200 Hz case, we chose the tone SPL to be higher (72 dB) in order to get similar levels of firing activity in the auditory nerve fibers as for the higher frequencies (Fig 2M, S1 Fig). In all cases, the background noise was chosen to have the same SPL level as the tone. As a result, the superposition of the tone and background noise was at a higher dB (typically a *≈*3 dB increase) than the tone or background alone. The control case was thus at a slightly lower dB level than the other 4 cases considered. For all of our analyses, 10 different realizations of background noise were simulated for each background case and tone frequency, and the model simulation results presented were averaged across results from these realizations.

**Fig 2.**
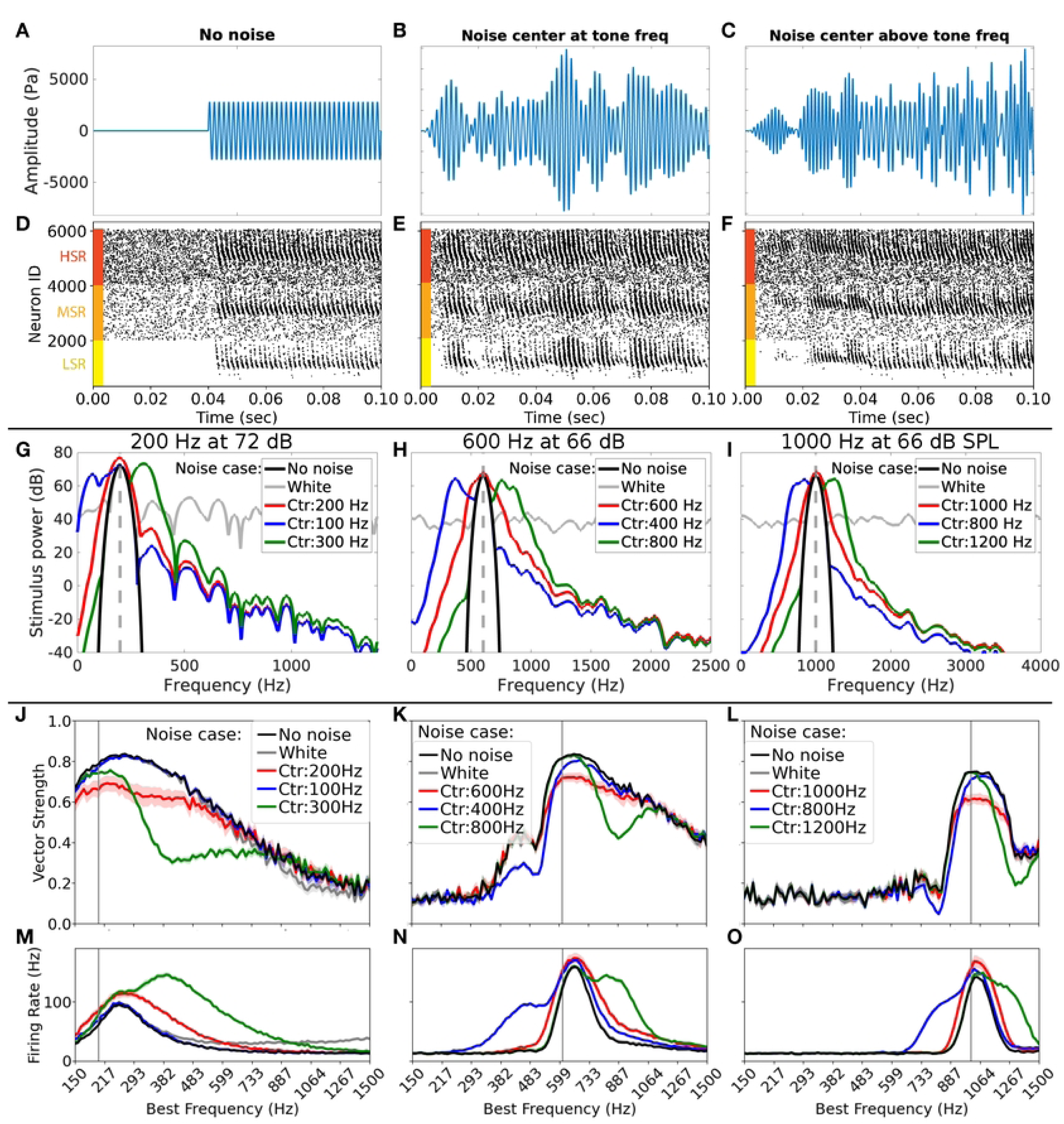
Types of sound inputs used and their impact on SGN spiking activity. A-C: Examples of input waveforms for the input tone frequency of 600 Hz. The input signal was a pure tone, with each simulated case consisting of 40 ms of background sound only, followed by 60 ms of the same background superimposed with the input signal. A: The pure tone (no noise) case, which served as the control. B: The centered noise case where the narrowband noise background had its central frequency at the tone frequency. C: Narrowband noise case with the center above the tone frequency. D-F: Example raster plots of SGN heminode spikes in response to input sounds depicted in A-C. Neurons were ordered in 3 sets for Low Spontaneous Rate (bottom third, indicated with the yellow bar), Medium Spontaneous Rate (middle third, orange bar), and High Spontaneous Rate (top third, red bar) classes. Within each class, neurons were ordered in increasing Best Frequency (BF) from 150 Hz to 1500 Hz. G-I: Stimulus power spectra for the 5 background noise cases chosen for each of the 3 tone frequencies considered (200 Hz, 600 Hz, 1000 Hz). The dashed gray vertical line denotes the tone frequency, the black curve is for the control case with no noise, the gray curve is for the case with added white noise, the red curve denotes the centered noise case, and blue and green curves denote the other two narrowband noise cases for each tone frequency. J-L: Vector Strength (VS) of SGN fiber responses with respect to the peaks of the corresponding sinusoidal tone (J: 200 Hz, K: 600 Hz, L: 1000 Hz); the X axis denotes BF of the fibers. VS was higher for BFs near input sound frequencies, and narrowband noise centered at tone frequency (red) led to the most degradation in VS at near-tone BFs, as compared to the no noise case (black) and other background cases (white noise in gray, and other narrowband cases in blue and green). M-O: Mean firing rate of SGN fibers vs BF for all noise cases shown in J-L. Fibers of a given BF were most active when input sound contained corresponding frequency components. For plots in each row, the Y axis is given in the leftmost plot.

### Peripheral auditory system model

To simulate neurotransmitter release dynamics at the synapses between inner hair cells (IHCs) and auditory nerve fibers, we used a previously developed model for the peripheral auditory system of guinea pig [23, 24]. In this model [23, 24], a second-order linear bandpass Butterworth filter with cutoffs of 22 kHz and 12.5 kHz is applied to the input signal to account for the response of the ear and compute the output stapes velocity. In response to stapes movement, the basilar membrane (BM) velocity is computed using a dual-resonance-nonlinear filter bank model, and 101 BM channels were simulated in this study. BM velocities are then used to approximate the displacement of IHC cilia, which are arranged tonotopically along the BM. The IHC cilia displacement changes the fraction of open ion channels at the IHC apical membrane, affecting the apical conductance which is modeled as a three-state Boltzmann distribution. The updated conductance leads to IHC membrane depolarization which is used to model the change in calcium current and thus calcium ion concentration near the synapse of the IHCs, affecting neurotransmitter release rate. The model for the IHC synapse is comprised of three pools of vesicles: a cleft pool, an immediate store, and a reprocessing store (refer to [23, 24] or [13] for all model equations and parameters). Vesicles in the cleft and reprocessing stores are continuous quantities, whereas the immediate store has quantal vesicles, whose release is a stochastic process. This process results in a release probability at time step *dt* of our input signal, which was sampled at 1*/dt*.

In the ear, IHCs synapse on the peripheral axon segments of the auditory nerves, Type I spiral ganglion neurons (SGNs, Fig 1A). The release probabilities output from the IHC-SGN synapse model were used to determine random times for synaptic release drawn from a uniform distribution. For every release time *t*, an external stimulus mimicking the post-synaptic current *I_app_* induced by release at the IHC-SGN synapse was applied to the model SGN axon:

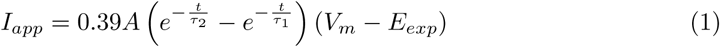

where *τ*_1_ = 0.1 ms, *τ*_2_ = 0.3 ms, *E_exp_* = 0 mV. The conductance of *I_app_*, *A*, was set to 0.14 nS to achieve appropriate SGN firing responses (see S2A,B Fig).

### SGN fiber model

A compartmental model of peripheral axons of Type I SGNs (referred to as SGN fibers throughout this study) was constructed using the NEURON simulator (version 7.6.2, [25]), depicted schematically in Fig 1B. The excitatory stimulus *I_app_* at the IHC-SGN synapse was input to the end of the first compartment, an unmyelinated segment of length *L_u_* which was connected to the heminode, the first node from the synapse. The position of this heminode was crucial in this study, as described in detail in Introduction. The heminode is followed by 5 myelinated segments separated by 4 nodes. Passive membrane properties for all compartments were defined by the specific capacitance (*C_m_*) and specific resistance (*R_m_*). Sodium and potassium channels were inserted along the SGN fiber except in myelinated segments, which have only passive membrane properties. The nominal conductances for ion channels along the initial unmyelinated segment were set to be 15 times less than at the nodes and heminode, and therefore, action potential generation occurs at the heminode. For details on the differential equations used in the model for membrane voltage and ion channel dynamics and the parameter values, refer to our previous work [13] and references therein. The differential equations were solved in the NEURON environment using a fully implicit backward Euler method with a time step of 5 *µ*s.

#### Defining fiber types

In line with our previous work [13, 14], we used three classes of SGN fibers defined by their spontaneous firing rates, thresholds for sound-evoked activity and saturation profiles: high spontaneous rate (HSR), medium spontaneous rate (MSR), and low spontaneous rate (LSR) fibers. To simulate these three different classes, we varied the maximal conductance of calcium ion channels and the minimum calcium concentration needed for neurotransmitter release in the peripheral auditory system model (see [13] for values) to mimic SGN firing responses reported in [26]. We incorporated fibers of 101 characteristic or best frequencies (BFs) ranging from 150 Hz to 1500 Hz, distributed according to the Greenwood function [24, 27]. 20 SGN fibers were simulated for each of the 3 spontaneous rate classes and the 101 BF classes, giving a total population size of 6060 SGN fibers. Each fiber received inputs from an individual IHC-SGN synapse, and spikes at the SGN heminodes were recorded for analysis and subsequent input to the SOC model.

#### Modeling myelinopathy in SGN fiber populations

To simulate the deviation in heminode positions observed in myelinopathy [12], we varied the length of the initial unmyelinated segment (*L_u_*) in the SGN fibers (Fig 3A), keeping all synaptic release probabilities and other fiber properties identical to the healthy model. For the healthy model, *L_u_* was set at 6 µm, for all 6060 fibers in the population. Two regimes of myelinopathy were considered: mild and severe, defined by the extent of variability in heminode positions. In the mild myelinopathy (MM) model, the population was divided into 4 equal-sized subpopulations with 1515 fibers each (each subpopulation contained fibers from all BF and spontaneous rate classes modeled in the healthy case). Fibers in the first quarter subpopulation maintained the healthy *L_u_* of 6 µm, while *L_u_* was increased beyond 6 µm in the remaining subpopulations. Specifically, in the second quarter *L_u_* = 8 µm, in the third quarter *L_u_* = 10 µm, and in the last quarter *L_u_* = 12 µm. In this MM regime, all the fibers, even those with the longest *L_u_* (12 µm), were able to spike normally in response to the simulated IHC-SGN synaptic input (see S2A,B Fig with the IHC-SGN synaptic strength of 14 nS). In the severe myelinopathy (SM) model, the SGN population was similarly divided into 4 equal quarters, but with a maximum *L_u_*= 14 µm, such that the four quarters of the population had *L_u_*= 6, 9, 12, and 14 µm. Here in the SM population, fibers with *L_u_* = 14 µm did not spike in response to excitatory input at the IHC-SGN synapse (S2A,B Fig).

**Fig 3.**
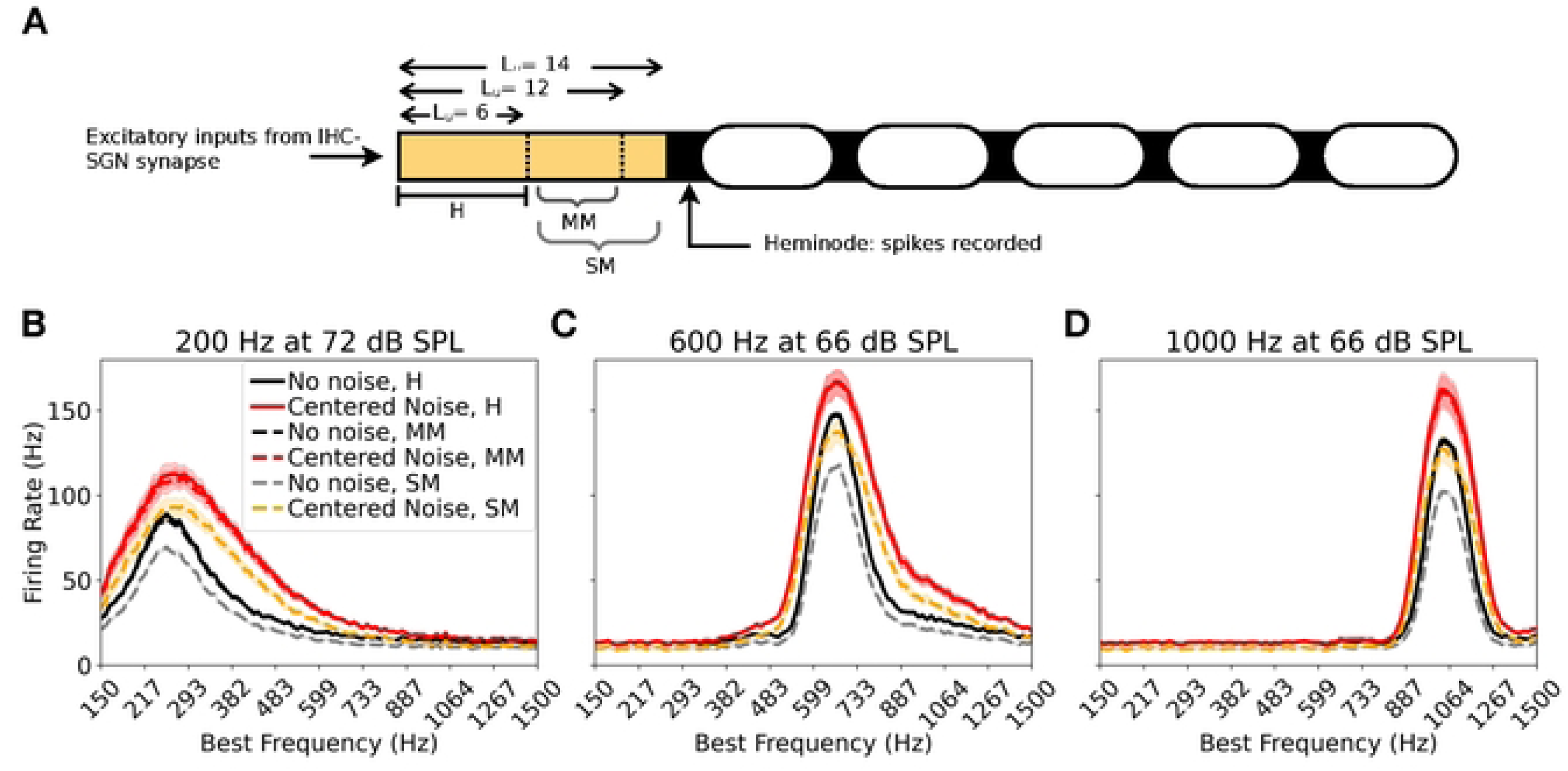
Introducing myelinopathy in the healthy model. A: Effects of myelinopathy were modeled by varying the length of the unmyelinated segment L_u_, which was 6 µm for all SGN fibers in the healthy (control) case (H). In the mild myelinopathy (MM) regime, the SGN population had fibers with unmyelinated fiber length (L_u_) between 6 and 12 µm, and in severe myelinopathy (SM), L_u_ ranged between 6 and 14 µm (see Modeling myelinopathy in SGN fiber populations,Effects of myelinopathy on SGN fiber activity patterns for details). B-D: Firing rate of SGN fibers as a function of best frequency (BF) for H, MM, and SM models shown for no noise and centered noise cases. For both background noise cases, MM (in dashed black and dashed red) showed similar firing activity as H (solid black and solid red lines, respectively) for all tone frequencies, while SM (dashed gray and dashed orange lines) showed *≈*25% fewer spikes than H and MM for all BFs. Y axis for B-D is shown in B on the left.

### Vector strength (VS) measurement

Vector strength (VS) is a measure of phase locking of a neuron population’s spiking activity with respect to the sound stimulus. To compute VS in response to a sinusoidal tone with well-defined peaks, we first computed the phase angle *θ_i_* of each spike *i* fired by cells in the population as:

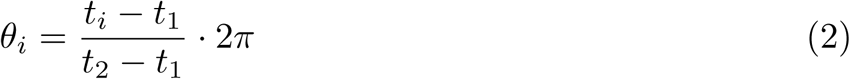

where *t_i_* is the time of the spike, and *t*_1_ and *t*_2_ mark the peaks of the sinusoidal stimulus just before and just after *t_i_*, respectively. VS was then computed as [28]:

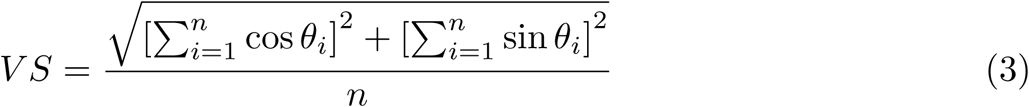

where *n* is the total number of spikes fired by the population. This measure varies between 0 and 1, where a value of 1 denotes perfect phase locking with every spike occurring at the same delay relative to the previous sound stimulus peak.

### Jitter analysis

To identify the source of any degradation to VS in myelinopathy or noisy conditions (see Results, Fig 2J-L and 4G-I), we assessed the variability in the spike times of SGN fibers in 2 ways. First, we looked at the jitter in spike times across the cycles of the sinusoidal pure tone sound signal. To do so, we computed the standard deviation of spike phases with respect to the sinusoidal tone peaks, considering all spikes after tone onset (at 40 ms, see Sound inputs) for each SGN fiber, and reported the average of this cycle-to-cycle jitter across SGN fibers of each BF class. Next, we considered the jitter in spike responses across the SGN fibers of each BF class. To do so, we computed the mean phase of all spikes after tone onset for each SGN fiber and computed the standard deviation of this mean spike phase across SGN fibers of a given BF. Circular statistics were used for this analysis, which is summarized again in Effects of myelinopathy on SGN fiber activity patterns and depicted schematically in Fig 4J.

**Fig 4.**
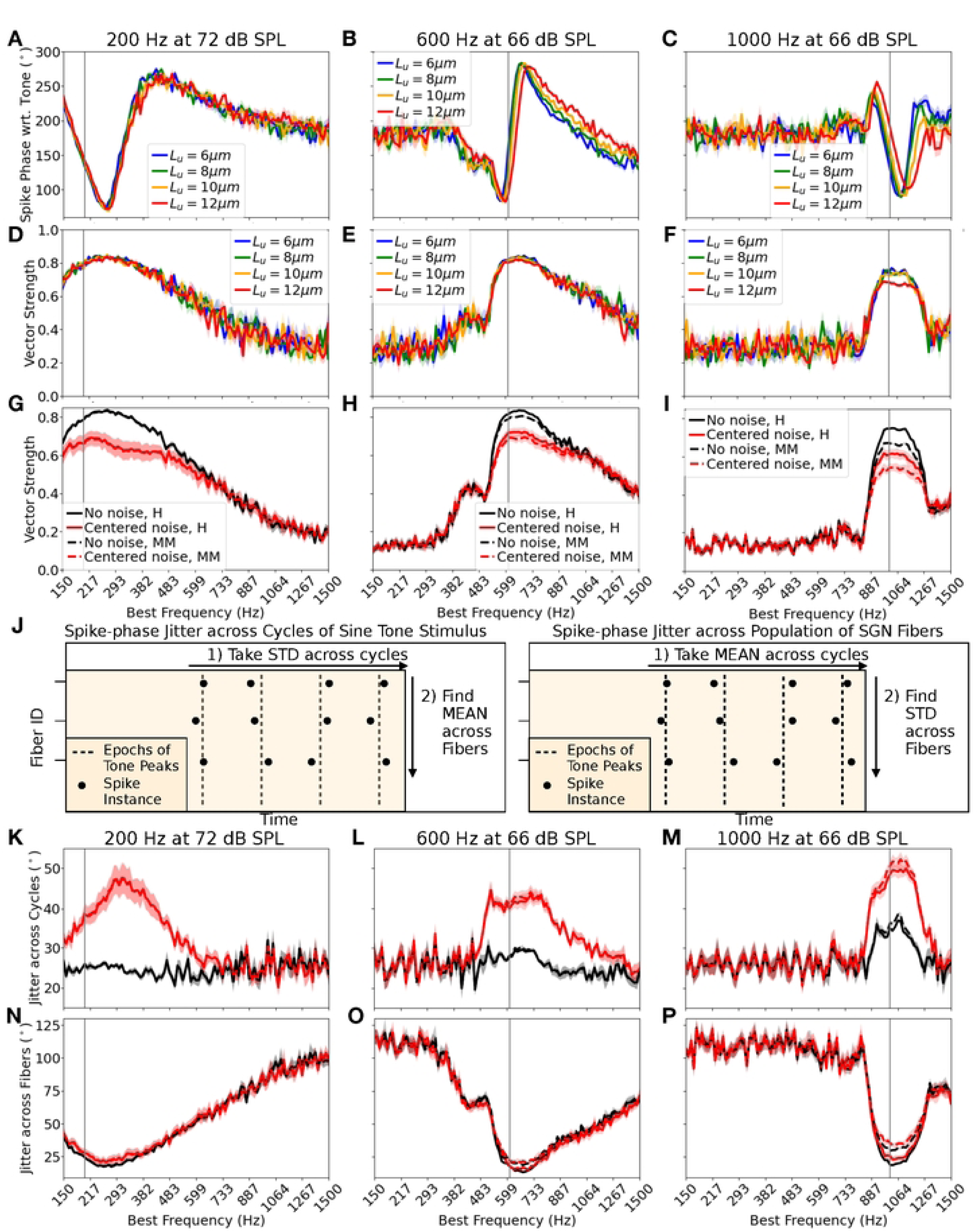
Frequency dependent impacts of mild myelinopathy on SGN firing activity. A-C: Effects of unmyelinated fiber length, L_u_, on mean phase of SGN spikes with respect to tone peaks, as a function of BF (X-axis), and for varying input tone frequency of A: 200 Hz, B: 600 Hz, and C: 1000 Hz. The input was a pure tone only. For higher frequencies, sub-populations of different L_u_ showed variable mean phases with respect to tone peaks, with the effect most pronounced for fibers with L_u_=12 µm. This trend was absent at 200 Hz, and most evident at 1000 Hz. D-F: Effects of unmyelinated fiber length, L_u_, on VS of SGN fibers, as a function of BF (X-axis), and for varying input tone frequency; D: 200 Hz, E: 600 Hz, and F: 1000 Hz. The input again was a pure tone. For 1000 Hz, the L_u_=12 µm sub-population showed clearly reduced VS. G-I: Cumulative effect of mild myelinopathy on SGN fiber VS as a function of BF, for the three tone frequencies; G: 200 Hz, H: 600 Hz, and I: 1000 Hz. Mild myelinopathy regime (dashed line) was compared with the healthy (solid line) case for no noise and centered noise input cases. Centered noise (red) consistently resulted in reduced VS for all three tone frequency cases, as seen earlier. Myelinopathy led to a reduction in VS in a frequency-dependent manner, with effects absent at 200 Hz (solid lines overlap with dashed lines), and most pronounced at 1000 Hz. J: Depiction of the measurement of two separate sources of spike variability in SGNs. Left: **Spike jitter driven by background noise:** we measured jitter in SGN spike-times across the peaks in the sinusoidal tone by recording the variance of spike phase with respect to tone peaks for all cycles during 60 ms of tone input and finding the mean variance across all fibers. Right: **Spike jitter driven by myelinopathy:** we measured jitter in SGN responses across fibers in the population by recording the mean spike phase over all cycles, and finding the variance of the mean phase across the population. K-M: Jitter across the cycles for; K: 200 Hz, L: 600 Hz, and M: 1000 Hz. For all three tone frequency cases, centered noise consistently increased jitter across tone cycles, explaining the reduced VS with noise in G-I. N-P: Jitter across the fibers for; N: 200 Hz, O: 600 Hz, and P: 1000 Hz. Myelinopathy increased jitter across fibers for higher frequencies only, with no change at 200 Hz (solid lines overlap with dashed lines), following the trend observed in G-I. For plots in each row, the Y axis is given in the leftmost plot.

### SOC model

SGN fiber activity is relayed to the MSO through SBCs and GBCs in the anteroventral cochlear nucleus. Specifically, multiple SGNs make excitatory projections to individual SBCs and GBCs on the ipsilateral side (2-4 to SBCs and 9-69 to GBCs [29–32]). MSO cells receive binaural excitatory inputs directly from SBCs and binaural inhibitory inputs from GBCs via relay points, the medial nucleus of the trapezoid body (MNTB, from the contralateral side) and the lateral nucleus of the trapezoid body (LNTB, from the ipsilateral side). Since MNTB cells act as accurate biological relays to the MSO [33, 34], and not much is known about the properties of LNTB cells [35–37], we chose not to include these structures and treated inhibitory MSO inputs as coming directly from GBCs (see [14]), in line with previous work [38]. Our reduced SOC model thus consisted of ipsilateral and contralateral SBC, GBC, and MSO cell populations, with each population containing 300 neurons (Fig 1C). Each SBC and GBC received 4 and 40 excitatory inputs, respectively, from ipsilateral SGN fibers. At the MSO, the first point of binaural integration along the auditory pathway, each cell received excitatory inputs from 6 SBCs on each side and inhibitory inputs from 3 GBCs on each side. GBCs and MSO cells were modeled as single-compartment Hodgkin-Huxley units with parameters and ion channel dynamics taken from [38] (also described in [14]), adjusted to closely resemble experimental results. Since both SBCs and GBCs have comparable dynamics, only varying in morphology and connectivity with SGNs [39], SBCs were modeled using the same neural dynamics as GBCs. The only change to this model from our previous model [14], was to the strength of the excitatory synapse between an SGN fiber and an SBC cell (*A* in Eq. 18 from [14]), which was increased by a factor of 1.2 to elicit a sufficient number of MSO cell spikes for analysis for all stimulus and myelinopathy cases simulated, including the SM models with very low spiking activity.

The auditory system is organized tonotopically, from the cochlea [40] through the cochlear nucleus and SOC [41, 42] and all the way to the auditory cortex [43]. This means that the various frequency components in input sounds are processed in parallel frequency channels, without much interference between these channels. Since we are studying how the auditory processing of pure tones is impacted in noisy settings and with myelinopathy, we have focused on localization results for the frequency band of interest (given by the tone). Thus, the spiking activity of only the SGN fibers with BFs near the tone frequency that showed high VS values with the pure tone was propagated to the SOC model for subsequent processing. For all tone frequencies and cases, 200 SGN fibers with near-tone BFs were selected.

## Results

Using our computational model, we investigated the effects of background noise, combined with the desynchronizing effects of myelinopathy on auditory coding, on sound detection and localization. We employed different sound stimuli with varying noise content. As described in the Methods section, each 100 ms sound waveform consisted of an initial 40 ms of background noise followed by 60 ms of a pure sinusoidal tone (at 200, 600, or 1000 Hz) superimposed on the background. We considered 5 cases for the background: no noise, white noise, and cases with narrowband filtered noise centered at frequencies matching the tone or above or below it (Fig 2A-C show some example waveforms for a tone frequency of 600 Hz).

### Effects of background noise on SGN population response activity

Using the healthy model for Type I Spiral Ganglion Neuron (SGN) fibers (see Materials and methods), we assessed the response of the SGN fiber population to the different sound stimuli. Fig 2D-F show raster plots of SGN fiber spiking activity in response to the sound waveforms shown in Fig 2A-C, respectively. In each raster plot, SGN fibers are grouped according to their spontaneous rate class (from top to bottom: high (HSR, indicated by red bar), medium (MSR, orange bar), and low (LSR, yellow bar) spontaneous rate), and sorted according to their best frequency (BF) (from 150 to 1500 Hz in ascending order along the Y axis) within each spontaneous rate class. In the no noise case, SGN fibers showed a clear increase in spiking above baseline firing when the tone starts (at 40 ms, Fig 2D) and spikes appeared to be phase-locked with peaks of the tone. As expected, only the fibers with BFs close to the tone frequency (600 Hz in this case) showed distinct activation patterns, with a clear difference in baseline and sound-evoked activity levels across the spontaneous rate classes.

For the centered noise sound waveform (Fig 2B), fibers of the corresponding BFs showed increased spiking even before the tone starts (first 40 ms, in Fig 2E). During the tone (after 40 ms), episodes of phase-locked firing were interspersed with periods of near-baseline activity, corresponding to intervals of constructive or destructive interference between the sinusoidal tone and noise frequency components near the tone. On the other hand, for noise centered above the tone frequency (noise center at 800 Hz in this case, Fig 2C), fibers with BFs near 800 Hz transiently responded to the background noise (Fig 2F), while the fibers with BFs near the tone frequency (600 Hz) continued to show control-like phase-locked activity in response to the tone.

The encoding of stimulus identity in the activity of the SGN fiber population was assessed through the extent of phase locking of SGN fiber spike trains with respect to the peaks of the pure sinusoidal tone. Disruption in encoding of the tone stimulus due to background noise was quantified using the Vector Strength (VS, see Vector strength (VS) measurement) computed for the activity of fibers of each BF. VS was computed relative to the tone using spikes generated only after the tone started, i.e., after 40 ms. For this analysis (in Fig 2J-O), spikes of SGN fibers from all 3 spontaneous rate classes were pooled together for each BF simulated. LSR fibers showed very low levels of activation, and so HSR and MSR fibers primarily contributed to the trends shown here (see S3 and S4 Figs). For the control case for all 3 tone frequencies, the VS peaks for fibers with BF close to the tone frequency approached 1 and those fibers exhibited the highest firing rates (black curves in Fig 2J and M for 200 Hz; K and N for 600 Hz and L and O for 1000 Hz) reflecting accurate encoding of the tone. For the 200 Hz and 600 Hz tones, VS was also high for fibers with BF above the tone frequency, indicating that these fibers also showed some phase-locking to the tone. With centered narrowband background noise (red curves in Fig 2J-O), we observed a remarkable decrease in the peak height of the VS levels for all frequency cases, indicating a reduction in phase-locked firing of near-tone BFs. Firing rates of fibers with BFs near the tone increased slightly due to the slightly higher dB level of the combined tone plus noise signal (see Sound inputs). For the other narrowband noise center cases (blue and green curves in Fig 2J-O), near-tone BFs showed slightly decreased VS levels, but not as much as with centered noise. However, fibers with BF near the noise centers displayed increased firing rates but decreased VS levels compared to control, indicating that the background noise disrupted their phase-locked firing to the tone. For all frequency cases, white noise background (gray in Fig 2J-O) led to no observable change in VS or firing rate compared to the no noise case (gray and black curves overlap), emphasizing the fact that the signal-to-noise ratio (SNR) at the tone frequency is the most critical feature of the background noise in disrupting tone encoding.

For subsequent analyses, we focused on the effects of the centered narrowband noise case in comparison with the pure tone case with no noise. This is motivated by the presumed minimal interference between SGN fibers and SOC neurons with different BFs due to the tonotopic organization of the parts of the auditory processing system accounted for in the model (see SOC model). Additionally, as the above results showed, noise power at the tone frequency most strongly affected the peripheral encoding of the tone. Thus, the centered noise case, which had the lowest SNR in the tone frequency band, was the most pertinent condition to analyze.

### Effects of myelinopathy on SGN fiber activity patterns

We altered our healthy peripheral model to simulate myelinopathy and analyzed deviations in the responses of the SGN fiber population to the sound stimulus. To simulate the observed variability in SGN heminode position in myelinopathy [12], we varied the length of the initial unmyelinated segment (*L_u_*) in the SGN fiber models (Fig 3A), keeping all synaptic release probabilities and other fiber properties the same as in the healthy model. For the healthy model, *L_u_* was set to 6 µm, for all 6060 fibers in the population. Two regimes of myelinopathy were considered: mild and severe, depending on the extent of variability in heminode positions. In the mild myelinopathy (MM) model, the population was divided into 4 equal-sized sub-populations with *L_u_* = 6, 8, 10, and 12 µm (see Modeling myelinopathy in SGN fiber populations). In this MM regime, the total number of spikes recorded at the heminodes for the entire population was similar to the healthy model for all frequency cases (Fig 3B-D). This is true for both no noise (compare solid and dashed black curves in Fig 3B-D) and centered noise (compare solid and dashed red curves in Fig 3B-D) cases. In the severe myelinopathy (SM) model, the *L_u_* lengths in the 4 subpopulations were *L_u_* = 6, 9, 12, and 14 µm. Due to the fibers with *L_u_* = 14 µm, the total number of spikes recorded at the heminodes for the entire population was about 25% lower than for the healthy model for all frequency values and both noise cases (compare solid black and dashed gray curves, and solid red and dashed orange curves in Fig 3B-D). It is worth noting that the number of spikes for the centered noise case was always higher than without noise due to a higher dB level (see Sound inputs).

We next investigated how myelinopathy combined with noise affected tone encoding in SGN fibers. Since we are interested in the temporal precision of SGN fiber spike timings with respect to the stimulus, we considered only MM for this analysis, where the model generated the same number of spikes as in the healthy case. We expect that the SM regime will display all the temporal disruptions seen in the MM case, but with additional degradation due to spike loss. Before analyzing population-level behavior with myelinopathy, we first looked at how each fiber subpopulation in MM (defined by *L_u_* = 6, 8, 10, 12 µm) responded individually to sound stimuli. We computed the mean value of the spike phase with respect to the preceding peak of the sinusoidal tone stimulus for fibers of all BFs (Fig 4A-C). For 200 Hz tone frequency (Fig 4A), all four *L_u_* subpopulations showed similar spike phases, meaning that irrespective of the unmyelinated segment length, the SGN fibers spiked with the same delay after a peak of the tone stimulus. However, for the 600 Hz tone (Fig 4B) and more prominently for the 1000 Hz tone (Fig 4C), the subpopulations with longer *L_u_* values (yellow and red curves) showed phase differences compared to the healthy fibers (blue curves). There is thus a frequency-dependent effect to be observed here, where the length of the unmyelinated segment preferentially affected SGN spiking patterns in response to higher frequencies. Vector strength values exhibited a similar frequency-dependent effect (Fig 4D-F): for only the 1000 Hz tone (Fig 4D), the VS of the longest *L_u_* subpopulation was degraded compared to the rest, while VS differences were not observed for 200 Hz and 600 Hz tones (Fig 4D, E). It is reasonable to expect the *L_u_*= 12 µm fibers to show more pronounced differences in spike timing compared to the shorter *L_u_* fibers due to the exponential and asymptotic nature of the delay in heminode spike timing with longer *L_u_* values (see S2A Fig, synaptic input conductance of 0.14 nS). However, it is interesting to note the frequency-dependent trend, where the impacts of the long *L_u_* perturbation are absent at lower frequencies.

Since with higher tone frequency fibers of different *L_u_* showed different mean spike phases, and also the longest *L_u_* subpopulation showed a reduced VS, we expected similar frequency-dependent trends to occur when we considered combined activity over all fiber subpopulations in the MM model population. Indeed, the degradation in VS in the MM condition was absent at 200 Hz (Fig 4G), but was present at 600 Hz (Fig 4H) and was pronounced at 1000 Hz (Fig 4I), for both the no noise (compare solid vs dashed black curves) and centered noise (compare solid vs dashed red curves) cases. Centered noise consistently degraded VS in healthy and MM models for all tone frequency cases (compare black vs red curves). To further analyze the source of spike-time variability leading to decreased VS, we analyzed the jitter in spike phases in two ways (see Jitter analysis for details): average jitter across cycles (Fig 4J, left panel) and average jitter across fibers (Fig 4J, right panel). Average jitter across cycles was computed (using circular statistics) as the standard deviation of the spike phase for all spikes after tone onset on each fiber, averaged across all fibers in the population. Average jitter across fibers was computed (using circular statistics) as the standard deviation across all fibers in the population of the mean phases of all spikes on each fiber after tone onset. We found that centered noise consistently increased average jitter across cycles for all frequency cases (Fig 4K-M, compare red vs black curves), explaining the VS degradation with centered noise (Fig 4G-I). Considering average jitter across the fibers (Fig 4N-P), for the 200 Hz tone, there was no clear separation between the noise or myelinopathy cases, but for higher tone frequencies, the MM model showed a higher jitter than the healthy model for both no noise and centered noise cases (compare solid vs dashed curves). These trends can explain the frequency-dependent effects on VS with MM in Fig 4G-I.

In summary, both mild levels of myelinopathy and near-tone frequency noise disrupted SGN encoding of the tone stimulus by reducing the ability of the SGN fibers to phase-lock with peaks in the tone waveform. Near-tone frequency noise had a consistent effect across all tone frequencies, while mild myelinopathy was specifically disruptive to high frequency tone encoding.

### Horizontal localization performance with noise and myelinopathy

Our previous modeling results [14] support the hypothesis that the observed peripheral auditory deficits in animal models of HHL can lead to perceptual deficits, by directly impacting the ability to localize sound cues in the horizontal plane. To analyze localization performance in the presence of background noise with varying levels of myelinopathy, we used a model of the mammalian superior olivary complex (SOC) circuit that includes cochlear sound processing and SGN, spherical and globular bushy cell (SBC and GBC), and medial superior olive (MSO) cell populations (see Fig 1C, SOC model, [14, 38]). MSO cell populations facilitate horizontal sound localization by acting as coincidence detectors, with their population firing activity showing sensitivity to interaural time differences (ITD) [44] (the time difference between sound waves arriving at the two ears). Individual MSO cells in a given hemisphere fire preferentially when they receive binaural excitatory inputs from the ipsilateral and contralateral SBC populations arriving at the same time. There is an intrinsic delay between neural inputs to MSO cells from the contralateral compared to the ipsilateral cochlear nuclei. As a result, the MSO population in a given hemisphere shows maximal firing activity when the sound source is slightly contralateral to that hemisphere so that the external and internal delays compensate each other. Thus, the MSO populations in the left and right hemispheres show bell-shaped response profiles as a function of ITD that are slightly displaced from each other, with the peak response of each hemisphere on opposite sides of the straight ahead position (ITD=0) (see S5A Fig). Thus, the difference in their firing rates, ΔR, varies with ITD (see S5B Fig). Generally, acuity for determining sound locations using ITD cues is best in the frontal hemifield rather than in lateral locations [45, 46]. For all of our analyses, therefore, we truncated the MSO ΔR response profile to a range of ITDs from −0.35 to +0.35 ms, which gives a roughly linear variation of ΔR with ITD (where ΔR=0 at ITD=0, see S5C Fig). Localization performance was assessed as the deviation of the ΔR profile from a horizontal line (no sensitivity to ITD), where steeper slopes of the ΔR curve generally indicate better localization performance. To model different ITDs for the tone stimulus, it was assumed to arrive at the left ear at phase 0 and to arrive at the right ear at phase = 2*π · ITD · f* for each tone frequency *f*. The background noise was assumed to be the same for both left and right ears for all tone ITDs to simulate a stationary background, but different horizontal locations of the source of the pertinent sound (see S5D Fig). For each ITD, SGN spikes were recorded for the left and right sides, and 200 SGN fibers with near-tone BFs were selected as input to the SOC circuit model (see Materials and methods for details).

We first analyzed localization performance of the MM and SM models compared to the healthy model in the control case (no noise). With MM, while the differences in left and right hemisphere MSO spike rates were similar to the healthy case in response to the 200 Hz tone (Fig 5A), the differences were smaller for the 600 Hz and 1000 Hz tones (Fig 5B, C). Thus, even without losing any spikes at the peripheral level, the jittered spike timing observed at the peripheral level could reduce horizontal localization performance. It is also worth noting here that localization performance was generally poor at 1000 Hz even for the healthy case (note difference in y-axis scale in Fig 5C); low frequency sounds only are localized at the MSO based on ITD cues, while higher frequency sounds are localized at the lateral superior olive (LSO) [47, 48]. For all tone frequencies, SM showed remarkably poorer performance compared with the healthy and MM cases. This indicates that the effect of the approximately 25% spike loss in the SM model was significantly exacerbated at the MSO.

**Fig 5.**
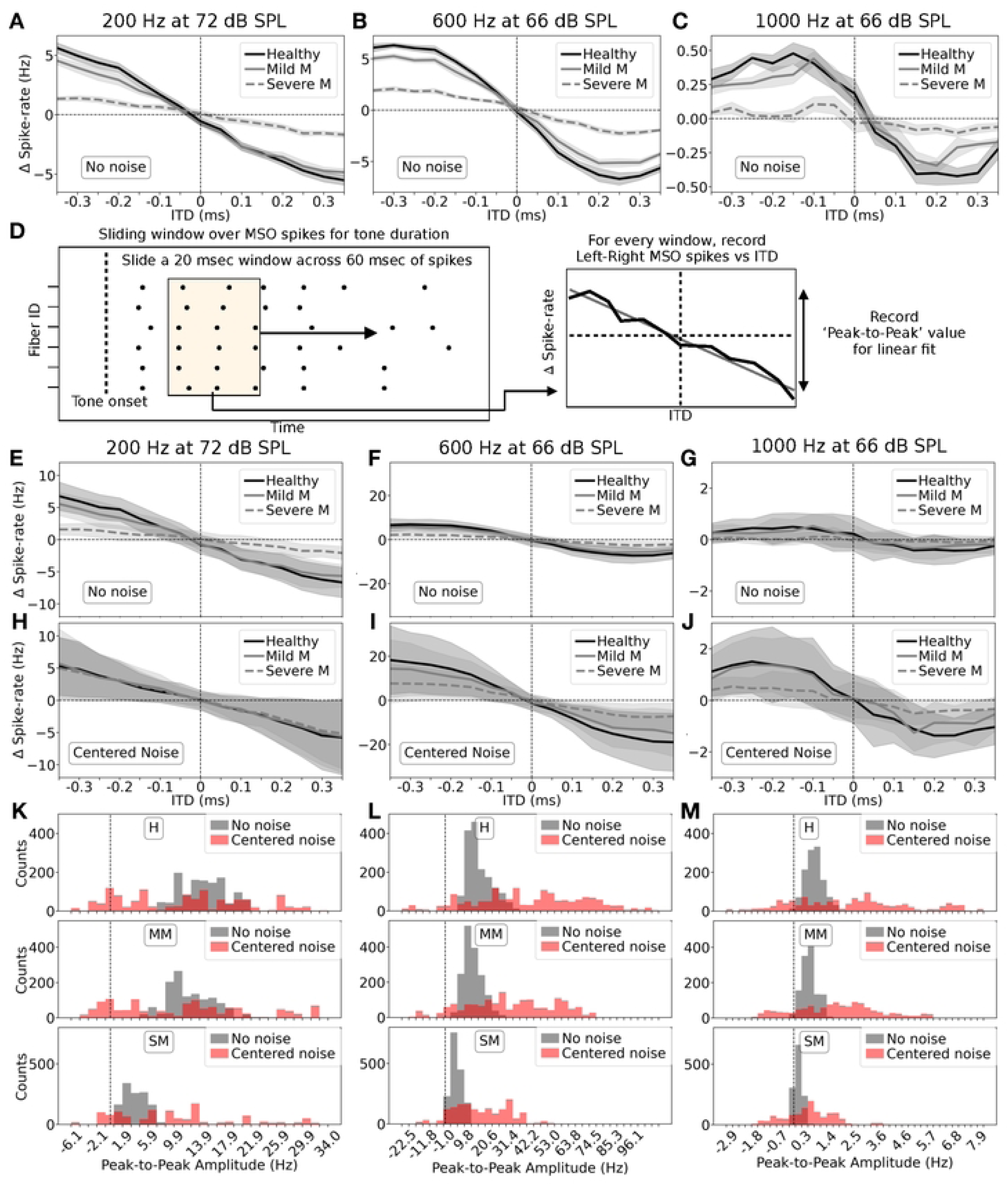
Localization performance at MSO with noise and myelinopathy. A-C: Difference in spike rate between left and right MSO populations for the entire input duration vs ITD, with no noise, and for the 3 tone frequencies: A: 200 Hz, B: 600 Hz, and C: 1000 Hz. Reduction in localization ability with mild myelinopathy (gray line) was more pronounced at higher frequencies than at 200 Hz, when compared with healthy localization (black line). Severe myelinopathy (dashed gray line) led to a remarkable reduction in localization ability for all 3 tone frequency cases. D: Analysis of localization performance across the temporal extent of stimuli: we slid a 20 ms window over MSO spikes corresponding to the input duration and measured binaural spike-rate difference for each window. The localization curve was recorded as a function of ITD. A linear fit was applied to the data for further analysis. E-G: No noise case; the mean and standard deviation over sliding windows (instead of the entire input duration as in A-C) of binaural spike rate difference is plotted as a function of ITD, for: E: 200 Hz, F: 600 Hz, and G: 1000 Hz input tone frequency. H-J: Same as E-G for the centered noise case. In all these plots, the standard deviations are large, more so for centered noise. This shows that there was a lot of variance in the localization profile over the input duration, making accurate localization harder. K-M: To study the changes in localization profile across the temporal extent of stimuli, we plotted histograms of the peak-to-peak value of the localization curves recorded over sliding windows (peak-to-peak value is the height of the localization curve: value at 0.35 ms ITD subtracted from the value at 0.35 ms ITD, shown in D) for different input signal frequencies (K: 200 Hz, L: 600 Hz, and M: 1000 Hz). For all cases and all tone frequencies, centered noise remarkably altered the localization profile of pure tones, with a large spread covering very high and very low (with some negative) peak-to-peak values. At the same time, mild myelinopathy biased both no noise and centered noise histograms towards the 0 peak-to-peak line (dashed vertical line), indicating degraded localization ability. This was even more pronounced for severe myelinopathy. For plots in each row, the Y-axis label is given in the leftmost plot.

For the results above, we considered the total number of MSO spikes fired in response to the input sound waveform (after the tone signal was turned on at 40 ms). However, we were also interested in understanding the variation in instantaneous localization across the duration of the sound waveform to understand how the random amplitude and phase modulation of the background noise disrupts MSO temporal processing. In general, a reduced slope of the localization curve given by Δ*R*, or large unexpected variations in Δ*R* at a fixed ITD, reflect a degradation in localization encoding. To analyze variation in instantaneous localization encoding, we considered MSO responses in 20 ms sliding time windows across the sound waveform (from 40 to 100 ms). For each ITD, all the left and right MSO spikes within each 20 ms time window were used to compute Δ*R* vs ITD profiles for that window (Fig 5D). The mean and standard deviation of these profiles were then computed for all time windows for each ITD value. Pooled results across 10 simulations are shown for the no noise (Fig 5E-G) and centered noise (Fig 5H-J) cases for the three tone frequencies. Δ*R* values exhibited variability across time windows for each ITD (shaded regions) in the healthy, MM, and SM models, which were larger with centered noise. Such variability in localization across the sound waveform duration, even without background noise and in healthy conditions, could be explained by the transient firing properties of MSO cells. Specifically, for a continuous pure tone stimulus, MSO firing rates reduced rapidly after the initial response at tone onset (see S6A,C Fig for an example raster plot). The presence of background noise exacerbated this MSO spike-rate variability. In particular, the sound waveform with narrowband noise centered at the tone frequency exhibited large amplitude peaks when the noise constructively interfered with the tone, interspersed with smaller peaks when the noise destructively interfered with the tone (see Fig 2B for an example with the 600 Hz tone case). As a result, MSO cells exhibited large fluctuations in firing rates across the waveform duration (see S6B Fig).

Thus, in addition to considering the mean Δ*R* profiles (Fig 5E-J), we considered the variability in localization performance across the sound signal duration. For the Δ*R* vs ITD profile computed at each 20 ms time window, we recorded the peak-to-peak value (the largest positive value at ITD=-0.35 ms minus the largest negative value at ITD=0.35 ms after considering a linear fit, Fig 5D) and plotted histograms of these values for all time windows and trials (Fig 5K-M). For all tone frequencies and noise and myelinopathy cases, the no noise histograms (gray bars) had right-skewed Gaussian-like shapes with a discernible center and width. This shape follows from the nature of MSO cells, which adapted their firing rate in response to continuous stimuli (see S6C Fig). It should be noted that the range of peak-to-peak amplitude values on the X axis is different for each frequency case, since even with a similar number of SGN spikes in each case, the SOC network dynamics vary with frequency (1000 Hz especially had very low values, as explained above). With centered noise (red bars), for all cases, the histograms lost their well-defined shape, showing a much larger spread in amplitudes with no discernible shape. We also find that in many centered noise cases, there are amplitudes with negative values, implying the Δ*R* profile for that window was flipped about the origin, namely sounds on the left would appear to come from the right and vice versa. Thus, even in healthy conditions, the presence of background noise completely altered the temporal localization profile for the tone stimulus computed at the MSO, potentially leading to sub-par or even incorrect computation of the horizontal location of the tone across its duration.

Myelinopathy had a consistent effect of degrading peak-to-peak amplitudes regardless of background noise. In MM (middle row), the amplitude values for all tone frequencies shifted closer to a peak-to-peak value of 0 (dashed vertical line, indicating no Δ*R* variation with ITD) compared to the healthy model (top row, more clearly for 1000 Hz, Fig 5M). This effect was much more pronounced with SM (bottom row). We note that for the 200 Hz centered noise case, SM did not show a reduction in peak-to-peak values (Fig 5H, K): the 200 Hz centered noise case with an average SPL of 75 dB led to robust MSO responses with SM, so H and MM cases did not fire noticeably more than the SM case, due to higher spiking probability in the auditory pathway when simulated at an SPL of 75 dB (see S7 Fig). Since peak-to-peak values closer to 0 denote poorer localization ability, MM and SM models generally predicted reduced localization encoding throughout the stimulus duration.

## Discussion

In this study, we extended our computational modeling of auditory processing to investigate how myelinopathy (specifically, heminode disorganization, as detailed in Introduction) at spiral ganglion neuron (SGN) fiber heminodes interacts with background noise to disrupt neural encoding and sound localization, potentially contributing to hidden hearing loss (HHL). Our previous work showed that myelinopathy contributes to peripheral sound processing deficits as well as degradation in sound localization performance through both disruptions in temporal responses, as well as loss of firing responses at the SGN fibers, and consequently at the cochlear nucleus cells and the MSO [13, 14]. Here, we studied myelinopathy-induced temporal distortions in firing activity with and without reduction in firing activity by modeling mild and severe regimes for myelinopathy, respectively. We found that even mild levels of myelinopathy without loss of firing responses could sufficiently perturb spike timings to create deficits in peripheral processing as well as sound localization, in a frequency-dependent manner. Moreover, in severe myelinopathy, localization performance was severely degraded across all frequencies studied due to reduced firing activity throughout the simulated auditory circuit. We also extended our previous work to include more realistic scenarios in noisy environments, and found that when the background noise had frequency content similar to the sound source, peripheral processing deficits induced by myelinopathy were exacerbated in an additive way. Further, the ability for horizontal source localization was degraded, and the computed source location fluctuated throughout the stimulus duration, presumably leading to difficulties in consistently localizing sound as the stimulus persists.

In our previous work, we simulated compound action potentials (CAPs) representing cumulative SGN fiber responses, effectively mimicking the ABR peak 1, for a pure sinusoidal tone at a high frequency of 10 kHz. Here, we focused on lower tone frequencies below 1000 Hz, where SGN firing patterns show robust phase-locking with incoming stimuli (see S8 Fig, which shows that the vector strength (VS) of the fibers falls for high pure tone frequencies). Analyzing the vector strength of SGN fiber responses with respect to peaks of sinusoidal tone inputs, we found that mild myelinopathy reduced the extent of phase-locking for higher simulated frequencies (600 and 1000 Hz, see Fig 4), but this effect was absent at lower frequencies (200 Hz, close to the fundamental frequency of human voice [49]). With severe myelinopathy, we expected the same disruption to be present, but with additional loss of SGN fiber spikes. Since ABRs are typically used to study responses to higher frequency tones [17, 18], we simulated CAPs for the 1000 Hz case and our two regimes of myelinopathy, following the same approach as in our previous work [13] (namely, convolving the cumulative SGN spike trains with a unitary response function; Fig 6A). We found that mild myelinopathy introduced a slight delay in the CAP without any changes in peak height, whereas severe myelinopathy additionally led to a noticeable change in peak height (Fig 6B), as observed in our previous results [12, 13]. Here, our analysis with SGN spike trains directly using VS may point to the need for new diagnostic methods that can study changes in such phase-locking rather than a cumulative far-field response, to detect mild myelinopathy effects, especially at lower frequencies.

**Fig 6.**
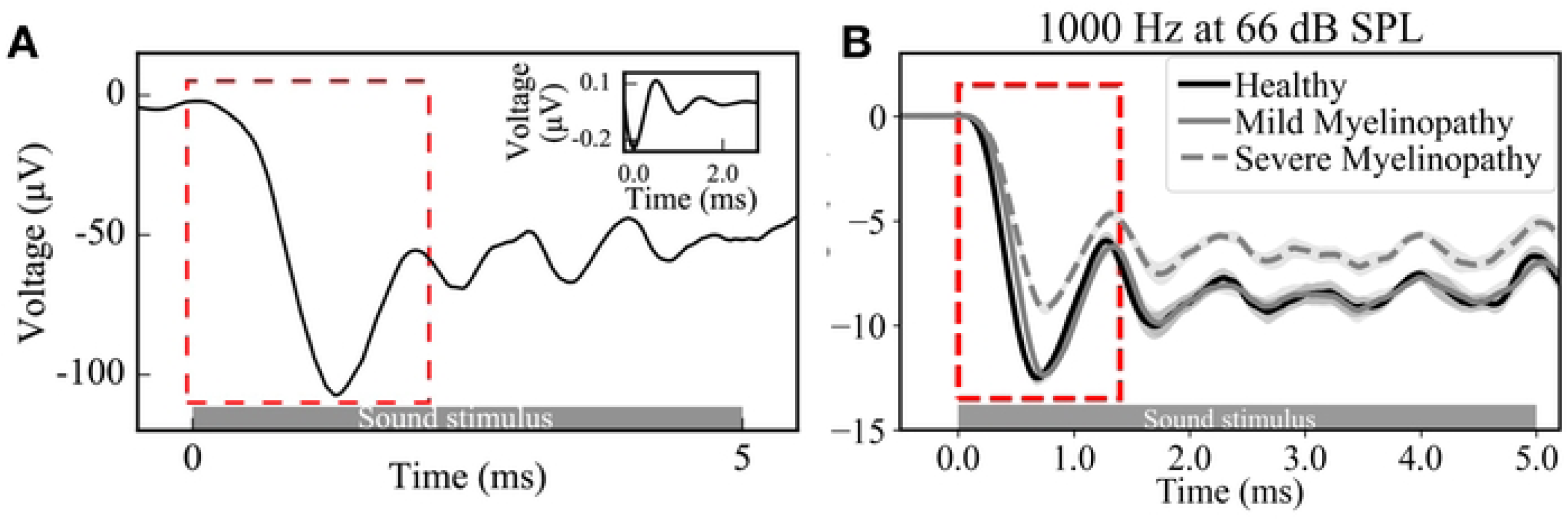
Simulated compound action potential (CAP) responses in different myelinopathy regimes. A: Example of a compound action potential (CAP) meant to simulate peak I of the Auditory Brainstem Response (ABR). Cumulative SGN firing activity was convolved with the unitary response function [50] shown in the inset, and the first negative peak was studied. B: CAPs simulated for healthy, mild, and severe myelinopathy cases for input tone frequencies of 1000 Hz. Mild myelinopathy regime showed a minor delay in the first CAP peak, with no change in peak height compared to the healthy case. Severe myelinopathy was necessary to produce a change in peak height that is clearly detectable in the ABR.

When we added background noise to the pure tone inputs, we found that only noise frequencies near the tone frequency disrupted the encoding of stimuli at SGN fibers, and that the reduction in VS due to noise and myelinopathy was additive, with noise reducing VS across all frequencies simulated (Figs 2,4). The reduction in VS is indicative of variability in spike phase with respect to peaks of the input tone, and our previous work has shown that any jitter in spike timing disrupts downstream processing by impairing the convergence of SGN inputs to cochlear nucleus cells [14]. We thus investigated the source of jitter by separately considering the jitter across the cycles of the sinusoidal tone for the stimulus duration, as well as the jitter in spike-timings between fibers in an SGN population (Fig 4). Model results showed that noise frequency components near the tone frequency increased the jitter in spike phase across the cycles of the input for all frequencies, which was expected since the background noise introduces a random phase shift to the input stimulus at any time instant. On the other hand, myelinopathy increased the jitter in phase-responses of fibers in the population, for higher frequencies only, explaining the frequency-dependent VS trends. These independent sources of jitter with noise and myelinopathy could also explain their additive effects seen in VS trends. Taken together, our results show that background noise can consistently induce jitter in peripheral auditory responses, which can be worsened by even mild levels of myelinopathy where firing responses are not lost, particularly for frequencies near 1000 Hz. Since controlling distinct degrees of myelinopathy is not possible to do in ablation experiments in animals, our results can provide unique insights into the mechanisms behind peripheral deficits observed in certain demyelinating diseases that may lead to HHL.

We have shown previously that high frequency tones (10 kHz) generate CAP responses with reduced peak height and increased latencies in severe myelinopathy scenarios consistent with experimental studies [12, 13], and consequently that jitter in spike timing coupled with loss in peripheral response rates lead to reduced localization ability [14] (considering a 200 Hz tone). Here, we provided a comprehensive analysis of the auditory localization circuit by using the peripheral responses for three tone frequencies (200, 600, and 1000 Hz) as inputs to the SOC model described in Materials and methods, and separately considering the mild and severe myelinopathy cases with and without spike loss, respectively. As expected from previous results, when myelinopathy was severe enough, such that spiking responses were lost throughout the auditory pathway, localization ability was severely affected across all frequencies. For mild myelinopathy, while localization performance was generally poor at 1000 Hz (MSO is specialized for localizing low frequency tones [47, 48]), we observed further reduction in the sensitivity to the horizontal location (Fig 5). Thus, model results suggest that perceptual deficits may occur with mild myelinopathy even when spike loss is not severe enough to be detected through peak height changes in ABRs. Furthermore, we observed that MSO firing responses showed transient behavior in response to sound tone, as they were modulated according to the background noise added (see S6 Fig). Therefore, we analyzed the evolution of the localization trends throughout the duration of the sound stimulus and compared the histogram profiles across myelinopathy cases with and without noise. Here, we could easily identify the effects of noise, which led to a large variability in localization sensitivity across input duration, corresponding to periods of constructive and destructive interference with the input pure tone (Fig 5). Additionally, the presence of mild myelinopathy biased all histograms towards lower localization sensitivities. Thus, presumably, these effects on localization sensitivity may contribute to speech intelligibility deficits reported in ‘cocktail party problem’ like scenarios [19]. In summary, our model results show that peripheral deficits introduced by noise and myelinopathy can lead to reduced localization ability, providing a possible link between peripheral deficits experimentally recorded in animals and human studies, and perceptual deficits in patients with myelin disorders [51].

Recent experimental studies analyzing the effect of noisy backgrounds on peripheral responses generated in response to human speech stimuli reported increasingly defective ABRs with increasing noise to signal ratios [15]. Our model could be extended to include realistic speech inputs and analyzed for different myelinopathy scenarios, as well as noise-to-signal ratios, to understand the firing pattern changes underlying these observed experimental results. Since our model simulates spiking activity in the auditory pathway up to the MSO, it could be tested on more complex input scenarios with a future goal to model multiple speakers by including two sinusoidal tones of different frequencies, and analyzing the localization profile for each tone in various myelinopathy and noise scenarios. A crucial shortcoming of the model is that it cannot account for responses in the SOC to higher frequency tones, which are localized in the Lateral Superior Olive (LSO) [47, 48], but extending our model to include the LSO could allow us to study higher frequencies, and directly link our findings to ABR changes observed experimentally. This would also allow us to use localization cues other than ITDs and expand the range of inputs that can be analyzed. Further, using transfer functions to simulate the modulation of spectra of incoming sounds by the external ear [52] could capture all cues necessary for localization.

## Conclusion

This study demonstrates that background noise and myelinopathy impair auditory processing through distinct but additive mechanisms that disrupt auditory processing at both the peripheral level and in subsequent localization networks. Our computational model showed that even mild myelinopathy (which preserved overall firing rates) degraded phase-locking of SGN fiber responses to pure tones in a frequency-dependent manner, particularly affecting the higher frequencies studied (600 Hz and above). By introducing variable phase shifts in the responses of affected SGN fibers, mild myelinopathy impaired the synchronous response of the auditory nerve in response to pure tone stimuli, eventually resulting in reduced localization ability in the MSO. We showed that added narrowband noise centered at the pure tone frequency further exacerbated these temporal deficits by introducing cycle-to-cycle variability in spike phases at SGN fibers, subsequently leading to fluctuations in sound localization sensitivity across the stimulus duration. This led to temporal instability at the MSO, where localization profiles could even invert during brief time windows in noisy conditions. In a severe myelinopathy scenario, where approximately 25% of peripheral spikes were lost, these effects were amplified throughout the auditory pathway due to reduced responsiveness at every level, resulting in profound localization deficits across all frequencies tested.

Our findings suggest that diagnostic approaches focusing on phase-locking measures using SGN fiber spiking patterns, rather than ABR peak heights that capture population-level SGN activity, may better detect mild myelinopathy at lower sound frequencies. By showing how heminode disruptions at the periphery contribute to localization deficits that are exacerbated in noisy conditions, our model results may offer explanations for scenarios like the “cocktail party problem” that affect speech intelligibility in humans. The model’s ability to isolate temporal distortions from spike loss provides insights into the physical mechanisms underlying different degrees of the pathology. Our biophysically detailed model separates the auditory processing pathway into distinct stages, which is helpful to understand the mechanisms underlying the processing of arbitrary stimuli, and allows for perturbations to the healthy model to capture any pathology of interest. Future extensions incorporating realistic speech stimuli, multiple simultaneous sound sources, and higher-frequency processing through the lateral superior olive would further extend our analysis to more complex real-world scenarios.

## Supporting information

**S1 Fig. Choice of SPL levels for frequencies considered.** The firing rate of SGN fibers in response to a pure sinusoidal tone is lower for lower tone frequencies at any given SPL. In order to obtain a similar spiking level in SGN fiber responses for all tone frequencies considered (to easily compare trends across frequencies), we chose a higher SPL for the 200 Hz tone (72 dB) than the 600 Hz and 1000 Hz tones (66 dB).

**S2 Fig. Choice of synaptic strength at IHC-SGN synapses.** A: Time between an excitatory input at the synapse and the first spike at the SGN heminode plotted as a function of L_u_ for various values of the synaptic strength A. All synaptic strength values showed exponentially increasing delays, with an asymptote at L_u_=14 µm for our choice of 0.14 nS (bold red). B: Probability of spiking at the SGN heminodes for various L_u_ and A values. At A = 0.14 nS (black rectangle), fibers with L_u_ 14 µm or more could not spike for any excitation at the synapse.

**S3 Fig. VS trends for SGN fibers of each spontaneous rate class.** A-C: VS of SGN fibers as a function of BF (X axis) of HSR fibers only, for all 5 background noise cases. D-F: VS trends of MSR fibers only. G-I: VS trends of LSR fibers only. LSR fibers showed very few spikes in response to sound stimuli, and our VS analysis using spike times resulted in noisy and unreliable trends. The VS trends that were calculated using spikes of fibers of all spontaneous rate classes pooled together (Fig 2J-L), thus mainly represent the firing activity of HSR and MSR fibers. J-R: VS trends for no noise and centered noise cases, with and without myelinopathy (J-L: HSR fibers, M-O: MSR fibers, P-R: LSR fibers). HSR and MSR fibers followed the same frequency dependent trend as seen in Fig4G-I, while LSR fibers generated too few spikes for meaningful analysis. Left column: 200 Hz tone case. Center column: 600 Hz tone case. Right column: 1000 Hz tone case.

**S4 Fig. Firing rate trends for SGN fibers of each spontaneous rate class.** A-C: Firing rate of SGN fibers as a function of BF (X axis) of HSR fibers only, for all 5 background noise cases. D-F: Firing rate trends of MSR fibers only. G-I: Firing rate trends of LSR fibers only. As expected, firing rate was highest for HSR fibers and lowest for LSR fibers. J-R: Firing rate trends for no noise and centered noise cases, with and without myelinopathy (J-L: HSR fibers, M-O: MSR fibers, P-R: LSR fibers). In all cases, mild myelinopathy led to no reduction in firing rate. Left column: 200 Hz tone case. Center column: 600 Hz tone case. Right column: 1000 Hz tone case.

**S5 Fig. Localization performance at MSO.** A: spike rate for left and right MSO populations for ITD values from −1 ms to 1 ms. Each side shows a peak in firing activity on opposite sides of the straight ahead position (ITD=0). B: The difference between left and right spike rates in A is sensitive to ITD. C: Within the frontal hemifield, this trend is roughly linear and can discriminate between ITDs to localize sound sources in the horizontal plane. D: Localization for input interval before 40 ms with centered noise only, showed no ITD dependence, as opposed to the interval containing the superimposed signal tone, post 40 ms. Only the signal tone’s location was modeled in a stationary background for all background cases.

**S6 Fig. MSO spike response analysis.** An example raster plot showing MSO spiking activity for the no noise case after the pure tone signal is turned on at 40 ms. Firing activity was highest at the onset of the tone, but adapted as the signal went on. B: With centered noise background, intervals of high activity were interspersed with quiet periods. C: Left panel shows three 20 ms time windows in the MSO spike rasters with no noise, during the input duration: one just after onset (left, in blue), and two windows in later stages (middle or mid in purple and right in red). Right panel shows localization curves as a function of ITD for these 3 windows, averaged over 10 trials. For later windows, the localization trend vs ITD became more horizontal, and this adaptation in MSO activity could explain the characteristic histogram shape seen in the no noise cases in Fig 5.

**S7 Fig. Comparison of MSO activity across frequencies and myelinopathy regimes for the centered noise case.** A: Spiking probability of SGN fibers in response to pure tone stimuli for different frequencies and dB levels, calculated as the firing rate in Hz divided by the corresponding frequency. With our choice of dB levels for the 3 frequency cases analyzed, the 200 Hz centered noise case with an average SPL of *≈*75 dB would result in higher spiking probability than the 600 Hz and 1000 Hz cases at *≈*69 dB, with similar effects at the MSO as well, as shown next. B-J: Example raster plots showing MSO activity with centered noise for the 3 frequency cases for the healthy case and the myelinopathy regimes. In the 200 Hz case, the healthy (B) and mild myelinopathy (E) cases did not fire significantly more than in the severe myelinopathy case (H), where most of the 300 MSO fibers simulated already responded robustly to peaks in the tone+noise waveform. However, in the 600 Hz and 1000 Hz cases (C,D: healthy, F,G: mild myelinopathy, and I,J: severe myelinopathy), severe myelinopathy showed a noticeable reduction in firing activity. This could explain the trends seen for the centered noise case at 200 Hz in Fig 5H and K.

**S8 Fig. SGN fiber phase locking regime.** VS of SGN fiber firing patterns with respect to peaks of a pure sinusoidal tone reduced with frequency, and showed high values of phase locking only for lower tone frequencies. We focused on this phase locking regime in this study.

## Acknowledgments

This work was supported by National Institute of Health grant NIH MH135565 (ST and MZ), R01DC000188 (GC) and R01DC018284 (MTR).

## Author Contributions

**Conceptualization:** Maral Budak, Gabriel Corfas, Victoria Booth, Michal Zochowski, Ross Maddox, Anahita H. Mehta, Michael T. Roberts.

**Formal analysis:** Siddhant Tripathy

**Investigation:** Siddhant Tripathy.

**Methodology:** Victoria Booth, Michal Zochowski.

**Software:** Siddhant Tripathy, Maral Budak.

**Supervision:** Victoria Booth, Michal Zochowski.

**Validation:** Siddhant Tripathy, Maral Budak.

**Visualization:** Siddhant Tripathy, Maral Budak, Victoria Booth, Michal Zochowski.

**Writing - original draft:** Siddhant Tripathy, Victoria Booth, Michal Zochowski.

**Writing - reviewing & editing:** Maral Budak, Ross Maddox, Anahita H. Mehta, Michael T. Roberts, Gabriel Corfas

